# The seed germination spectrum of 528 plant species: a global meta-regression in relation to temperature and water potential

**DOI:** 10.1101/2022.08.24.504107

**Authors:** Keyvan Maleki, Elias Soltani, Charlotte E. Seal, Hugh W. Pritchard, Jay Ram Lamichhane

## Abstract

The germination niche of plant species depends primarily on the seeds’ responsiveness to temperature and water potential. However, to appreciate future climate risks to natural regeneration through germination, a global level synthesis across species is needed. We performed a meta-regression of primary data from 377 studies on 528 species, including trees, grasses, crops and wild species, to determine patterns and co-correlants in the cardinal values that define species’ germination niche. A negative correlation was found between thermal time and base temperature, and positive correlations between other cardinal temperatures and base temperature. Mean values of thermal time indicate that annual crops germinate more rapidly compared to wild species, potentially as a consequence of domestication, and tropical tree seeds the slowest. Dryland species (Cactaceae and Agavaceae) have the widest upper thermal and lower moisture niche, indicative of an ability to grow under harsh conditions, while forages have the narrowest thermal and moisture niche, suggesting higher sensitivity to frost or drought. We propose a new conceptual framework for understanding germination niche as shaped by thermal and moisture traits. Our database represents a unique source of information to further determine the vegetation boundaries of wild or cultivated species, including within simulation studies on plant species adaptations under changing land-use and climate.

## Introduction

Germination is a critical growth stage both for domesticated and wild plants and, as such, it plays a vital role in their reproduction and regeneration. As the main environmental drivers, temperature and water potential modulate the germination response of all species (Baskin & Baskin, 2014). Water potential is the most important factor required for successful seed germination, influencing the vigour and final germination level/percentage. The soil is the main source of water for seeds from the environment (Bewley *et al*., 2012; Zhang *et al*., 2020). Temperature-regulated seed germination may act either by regulating dormancy status or by controlling the capacity and rates of germination (Bewley *et al*., 2012; Baskin & Baskin, 2014; Soltani *et al*., 2017a,b). In the presence of adequate moisture, the germination process is limited to a permissible range of temperatures that can be defined by ‘cardinal’ values; beyond these metabolic activities are impacted and germination does not progress (Bewley *et al*., 2012). A comprehensive understanding of seed germination behavior and description of intra- and inter-variabilities between groups of species (perennials and annuals, trees, grasses, crops and wild species etc.) to these two main environmental drivers of temperature and water potential is thus informative for two key reasons. First, in defining where seed-producing species are currently able to regenerate on the planet, and second, in predicting how these species will respond to unpredictable environments in the future (Fenner & Thompson, 2005; Baskin & Baskin, 2014; Gremer *et al*., 2020a,b). Moreover, environmental cues, as an indication of climatic factors, shape the germination niche of plant species that are responsive to specific germination requirements, including dormancy loss under warm and dry conditions, contributing to the timing of germination (Carta *et al*., 2022).

Identification of threshold-type responses to temperature and water potential values help characterize each species’ potential for regeneration via seed germination. Germination takes place between a minimum (hereafter referred to as base temperature; T_b_) and a maximum (T_max_) threshold temperature (also known as the ceiling temperature, T_c_), with the highest germination speed used to define the optimal temperature (T_opt_) (Bewley *et al*., 2012; Baskin & Baskin, 2014). T_b_ is defined as the predicted minimum temperature to be exceeded for germination to progress. It follows that germination will not progress at or below this temperature (Garcia-Huidobro *et al*., 1982a; Dahal & Bradford, 1994). Estimates of T_b_ are also valuable in calculating thermal times for the completion of germination in the sub-optimal range between the T_opt_ and T_b_. As thermal time varies with sub-populations/percentiles, such estimates represent a powerful means for predicting germination efficiency amongst a seed population under any, including changing, environmental conditions (Gummerson, 1986; Dahal & Bradford, 1994; Finch-Savage *et al*., 2005; Maleki *et al*., 2021). This concept of the thermal parameters of germination can be expanded by incorporating different water potentials, leading to hydrotime and hydrothermal time models describing the seed response above a base water potential for germination (b) and its interaction with temperature, respectively (Gummerson, 1986; Dahal & Bradford, 1994; Finch-Savage *et al*., 2005; Donohue *et al*., 2010; Bewley *et al*., 2012)

Estimates of thermal times, cardinal temperatures (i. e. T_b_, T_opt_ and T_max_) and _b_ have been quantified for many species, allowing an interpretation of seed germination within different environments (Trudgill *et al*., 2005; Orrù *et al*., 2012; Dürr *et al*., 2015; Seal *et al*., 2017; Zhang *et al*., 2020; Maleki *et al*., 2021) as well as intra- and inter-species comparisons facilitating prediction of their spatial distribution (Dürr *et al*., 2015). How crop and weed establishment occur under current and future climates (Forcella *et al*., 2000; Dürr *et al*., 2001; Gardarin *et al*., 2012; Lamichhane *et al*., 2020, 2022) and how plants synchronize germination and subsequent seedling growth with favorable conditions can be interpreted quantitatively (Dürr *et al*., 2015; Gremer *et al*., 2020a; Maleki *et al*., 2021). Application of these approaches for agricultural and ecological purposes has proven to be useful for an increased understanding of crop diversification in space (intercropping, relay cropping) and time (e.g. introduction of cover crops between two cash crops, double cropping). Designing innovative cropping systems as well as improving environmental sustainability (e.g. restoration of forests, dunes, arid lowlands) also benefit from thermal modelling (Waha *et al*., 2020; Fernández-Pascual *et al*., 2021; Beillouin *et al*., 2021).

A comprehensive review of cardinal temperatures for germination indicated that species originating from different geographical origins may show variable cardinal temperatures (Dürr *et al*., 2015), and this response may result from evolutionary adaptations (Donohue *et al*., 2010; Baskin & Baskin, 2014). How trait– environment interactions can shape the germination response through the thermal niche has been comprehensively modelled in the laboratory (Thompson & Ceriani, 2003; Catelotti *et al*., 2020) and *in situ* (Porceddu *et al*., 2013; Blandino *et al*., 2022). Thermal time models in combination with relevant environmental parameters have been widely used to investigate thermal niche and germination responses to accumulated temperature in agricultural and more natural settings (Garcia-Huidobro *et al*., 1982b; Covell *et al*., 1986; Pritchard & KR., 1990; Hardegree, 2006; Porceddu *et al*., 2013; Dantas *et al*., 2020; Maleki *et al*., 2021). Previous studies have shown that variation in threshold-type responses to ongoing environmental conditions among species may reflect the seeds’ thermal memory of the maternal environment (Fernández-Pascual *et al*., 2019), as evidenced by the cumulative thermal effects on seed development (Daws *et al*., 2004; Baskin & Baskin, 2014) and varying levels of seed dormancy (Pritchard *et al*., 1999; Porceddu *et al*., 2013).

The concept of ecological niche has been used to define the breadth of thermal and moisture ranges in which seeds should be able to germinate under current or future climatic conditions (Porceddu *et al*., 2013; Sultan, 2015; Catelotti *et al*., 2020; Ordoñez-Salanueva *et al*., 2021). While the thermal time approach has been used to predict the consequences of climate change (Orrù *et al*., 2012; Seal *et al*., 2017; Dantas *et al*., 2020), hydro time and hydrothermal time quantification should better enable predictions of the impact of global environmental change on species’ emergence and habitat patterns. Such insight can then contribute to the design of amelioration and adaptation strategies for environmental management based on seed fitness. A previous study proposed a qualitative conceptual framework to define the temperature tolerance ranges of plant species (Walck *et al*., 2011) while more recent studies have shown that tropical plants will face the greatest risk from climate warming as they experience temperatures closer to their upper germination limits (Seal et al., 2017; Sentinella *et al*., 2020). However, no large scale study has yet quantified the germination sensitivity to temperature and moisture that is useful to predict the potential impacts of shifts in environmental conditions on germination.

Meta-analysis, meta-regression and systematic review have been widely used to analyze and synthesize data from published and unpublished sources (Borenstein *et al*., 2021). By employing a simple regression analysis for synthesizing results from multiple studies, meta-regression seeks consensus on significant scientific issues (Borenstein *et al*., 2021). To this aim, we built on the framework used by Dürr et al. (2015) for a global analysis of >200 species’ seed germination response, but with two major differences. First, we updated their database, which included 223 articles previously published on plant species with worldwide distribution, by including data from an additional 154 studies published in the last decade on threshold-type responses to both temperature and water potential conditions. Second, while Durr et al. (2015) used a simple literature mining method, we applied a meta-regression approach and employed a new statistical method to interpret changes in threshold-type responses to temperature and water potential values and the relationships among the tested traits. To this aim, we first took seven classes of variables into account, including the variation in ranges of critical temperature and water potential values among species. We then assessed how each species has variable threshold-type values, and what the relationship among traits would be. The information we collated provides new insight into the relevant ecologically-meaningful traits required for plant recruitment and reproduction that must be considered in studies within a quantitative framework for spatial distribution of habitats, and species dispersal.

## Materials and method

### Data collection

We updated a previous database from Dürr et al. (2015) that was based on 223 articles published before December 2011. We integrated additional data that were extracted from 154 articles published between December 2011 and April 2021. We used ISI-Web of Science database to retrieve these articles by using the keywords “thermal requirements + germination” (199 publications), “cardinal temperature + germination” (194 publications), “germination + hydrotime” (94 publications), “germination + hydrothermal time” (275 publications), and “thermal ranges + germination” (429 publications) for a total of 377 publications. The original dataset can be accessed at https://doi.org/10.15454/XP3XHW (see Data Availability statement). We used Microsoft Excel (version 2016) to collate data on traits related to hydro-thermal times and germination thresholds from the published papers (Garcia-Huidobro *et al*., 1982b; Gummerson, 1986; Dahal & Bradford, 1994). The traits considered in this study included cardinal temperatures (T_b_, T_opt_ and T_max,_ which are estimated through regression procedure), thermal time required for germination (θ_50_, which is calculated via *θ*_sub_ (g) = (*T*-*T*_b_) *t*_g_ and *θ*_sup_ (*g*) = (*T*_c_ - *T*) *t*_g_), and base water potential (_b_Ψwhich is quantified by 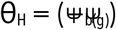 and 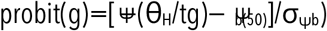. For further information, see Garcia-Huidobro *et al*., (1982), Gummerson (1986), Dahal & Bradford (1994) and Maleki et al. (2021), and references therein. For literature mining, seven categories of plant species were defined as previously (Dürr *et al*., 2015): crops, horticultural species (vegetables, leafy vegetables, ornamentals and medicinal plants), forage and rangeland species, Cactaceae and Agavaceae, wild species (invasive, endangered, wild potential medicinal plants), tropical trees and other trees. All perennial plants (trees and shrubs) were put into the trees category. To collect data on forage species, crops, and vegetables, only species that are important for agricultural purposes, were taken into account. Our study was premeditatedly limited to species with non-dormant seeds, or to seed lots where a pretreatment was applied to release dormancy to avoid the influence of differing patterns of dormancy on germination and on their interrelationships. Furthermore, weeds were excluded because of considerable variations in dormancy level as dormancy status can change cardinal temperatures (Baskin & Baskin, 2014).

### Determination of germination niche

We proposed a new conceptual framework based on the assumption that species construct their germination niche in response to environmental conditions they have experienced (Sultan, 2015; Fernández-Pascual *et al*., 2019). In considering germination niche, we took into account both thermal and moisture dependency. Then, we collected all available data on threshold-type responses to _b_Ψand cardinal temperatures. We then used a well-established framework for explaining the seed thermal niche. Based on the framework, thermal niche falls into two distinct categories, namely, the sub-optimal temperature range and the supra-optimal temperature range.

The sub-optimal temperature range, from T_b_ to T_opt_, in which the germination rate increases as the prevailing temperature rises up to T_opt_. The supra-optimal temperature range, from T_opt_ to T_max_, in which the germination rate becomes progressively slower with temperature increase. To determine differences in moisture niche shaped by _b_ values among plant categories, we set a range varying from 0 to negative values assigned to each plant category, suggesting a range in which seeds are able to germinate rapidly compared to the niche range occupied. To calculate both dimensions of niche, we computed the average potential and cardinal temperature values for each plant category, and then plotted the computed values on separate graphs to illustrate how the ecological niche might be distinct among species and plant category.

### Meta-regression analysis

Meta-regression is conceptually similar to simple linear regression, in which explanatory variables predict response variables. Here, the slopes of meta-regression are the effect size predicted by a regression line. Meta-regression coefficients explain how the response variable changes with an increase in the explanatory variable. The effect estimate denotes log risk ratio. Explanatory variables define aspects of studies that have considerable impact on the effect size (also called co-variates). A meta-regression method differs from linear regression in two ways: in a meta-regression, larger studies are more influential than smaller ones; and the residual heterogeneity can be modelled by explanatory variables, giving rise to random-effects meta-regression. The correlation between the tested traits is indicated by positive regression slopes with significant P-values.

To perform meta-regression analysis, the following standard regression model was used;

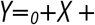

Where _*0*_ is intercept showing the overall effect size, *Y* denotes outcome variable that estimates the changes in traits of interest. *X* is matrix of explanatory variable. and indicate the vector of coefficients and the random error that refers to the classical regression model, respectively.

Weighted least-squares estimators of and 0 were computed as follows (Hartung *et al*., 2008);

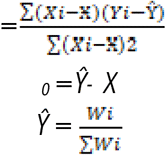

*Ŷ* denotes the estimates of the population effect size, *Wi* is weight of each study.

Residual sum of squares was calculated using following equation:

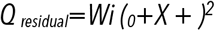

*Wi* is weight of each study.

We employed log risk ratio as dependent variable and threshold-type traits were used as co-variate variable;

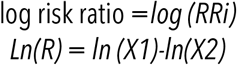

X1 is T_b_ values regressed on X2 representing other traits, including T_max_, T_oopt, b_ and _50_

The variance of log risk ratio (*vlnR*) is calculated as follows;

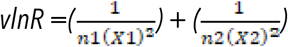

Where n_1_ and n_2_ denotes number of studies incorporated into analysis; X_1_ and X2 are T_b_ values regressed on X2 representing other traits, including T_max_, T_oopt, b_ and _50_

The approximate standard error (SE) was computed as follows:

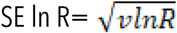

We performed meta-regression on only one covariate (e.g. T_b_ vs. T_max_), and, therefore, we suggested the possibility of the Z-test to examine its relationship with effect size. This meta-regression is thus based on the Z-distribution, which is a statistical approach to test the significance of the regression slopes. Therefore, we reported the Z-value with a corresponding p-value to indicate significant correlations, and, we also computed the magnitude of the relationship. The relationship of traits to effect size (defined as log risk ratio) is calculated as follows:

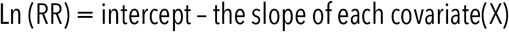

Where X is the absolute value of traits.

The 95% confidence interval for each covariate is estimated as follows:

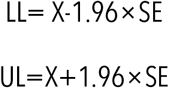

LL and UL refer to lower limit and upper limit, respectively. In the above-mentioned equations, 1.96 shows the Z-value corresponding to confidence limits of 95% (making space for 2.5% error at both end of the distribution).

## Results

### Intraspecific variation in traits among plant categories

Values related to intraspecific variation in traits among plant categories are reported in **Table 1** while the number of species included in intraspecific variation in the analyzed traits among the plant categories is presented in **Figure 1**.

**Table 1.** Intraspecific variation in the analyzed traits among the plant categories. The estimates were separately computed using the standard deviation of observed values for each category. T_b_, T_opt_ and T_max_, ΔT_opt_ -T_b_, ΔT_max_–T_opt_ indicate the base, optimum, maximum sub- and supra optimal temperature values, respectively; θ_50_ indicates thermal time required to attain 50% germination while _b_ indicates the base water potential values.

| Plant category | $T_b$ (C) | $T_{opt}$ (C) | $T_{max}$ (C) | $\Delta T_{opt}-T_b$ (C) | $\Delta T_{max}-T_{opt}$ (C) | $\theta_{50}$ ($^{\circ}\text{Cd}$) | $\psi_b$ (MPa) |
| --- | --- | --- | --- | --- | --- | --- | --- |
| Tropical trees | 4.034 | 3.404 | 4.852 | 5.352 | 6.006 | 134.113 | 0.392 |
| Other trees | 4.993 | 9.732 | 8.199 | 9.012 | 11.794 | 44.567 | 1.018 |
| Crops | 4.589 | 5.160 | 5.722 | 5.071 | 7.595 | 12.791 | 0.578 |
| Horticulture | 4.106 | 5.679 | 5.616 | 3.809 | 4.185 | 23.200 | 0.266 |
| Forage and rangeland | 2.322 | 4.599 | 4.735 | 3.456 | 5.037 | 12.495 | 0.344 |
| Cactaceae and Agavaceae | 3.477 | 4.504 | 5.535 | 4.527 | 5.366 | 19.434 | 0.569 |
| Wild species | 4.657 | 7.438 | 6.046 | 6.386 | 4.200 | 43.880 | 0.448 |
| Average | 4.025 | 5.788 | 5.815 | 5.373 | 6.343 | 41.497 | 0.516 |

**Figure 1.**
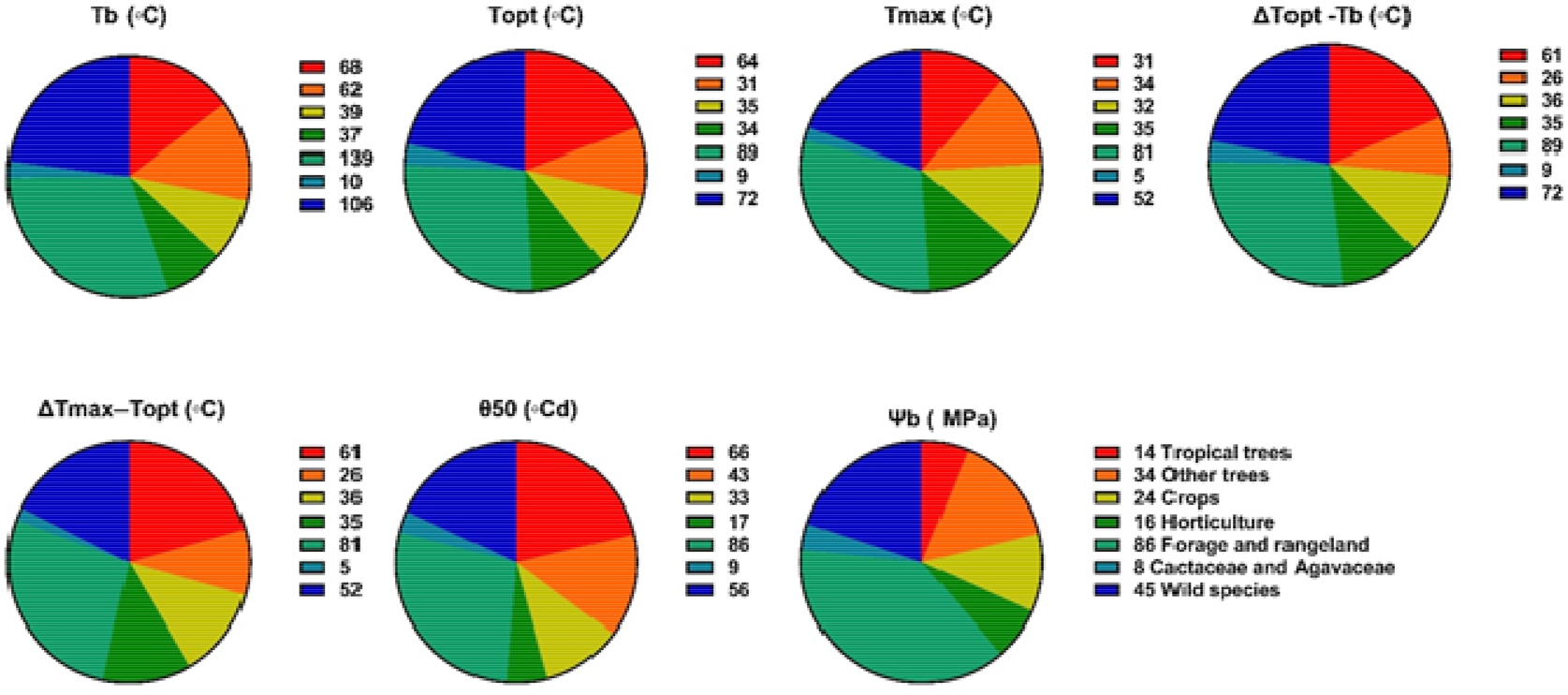
Number of species included in intraspecific variation in the analyzed traits among the plant categories. The numbers reported in the legend show the number of species in this set of analysis (the number of studies is provided in parenthesis of Figure 1). The number of species exceeds the number of studies as a given study may have reported data on different species. T_b_, T_opt_ and T_max_, ΔT_opt_ -T_b_, ΔT_max_–T_opt_ indicate the base, optimum, maximum sub- and supra optimal temperature values, respectively; θ_50_ indicates thermal time required to attain 50% germination while _b_ indicates the base water potential values.

Much information was available on forage and rangeland (198 species) and wild species (107 species) followed by tropical trees (85 species) and crops (45 species). In contrast, we found much less data on Cactaceae and Agavaceae (10 species). The range of variation in T_b_ was similar among plant categories with the ranges varying from 3°C for forage and rangeland to nearly 5°C for other trees. There was a considerable variation in _50_ with a window ranging from 12°Cd in forage and rangeland to 134°Cd in tropical trees. Variation in T_opt_, T_max_ and ΔT_opt_ - T_b_ was not important among plant categories. _b_ values showed the lowest variation among the plant categories. Among all traits considered, forage and rangeland species included higher number of species followed by wild species and tropical trees **(Figure 1)**.

### Cardinal temperatures

While Ψ_b_ values were reported only for 226 species, temperature-based traits were the most studied with T_b_ and T_opt_ values being reported for 461 and 334 species, respectively. T_b_ values varied greatly among plant groups, ranging from -4.8°C for *Koeleria vurilochensis* (i.e. forage and rangeland category) to 21.9°C for *Terminalia brassii* (Tropical trees category; **Figure 2a**). Succluent species (Cactaceae and Agavaceae families) showed less extreme values with the same mean of 10 C while other species groupings in the analysis had wider variabilities in T_b_ values.

**Figure 2.**
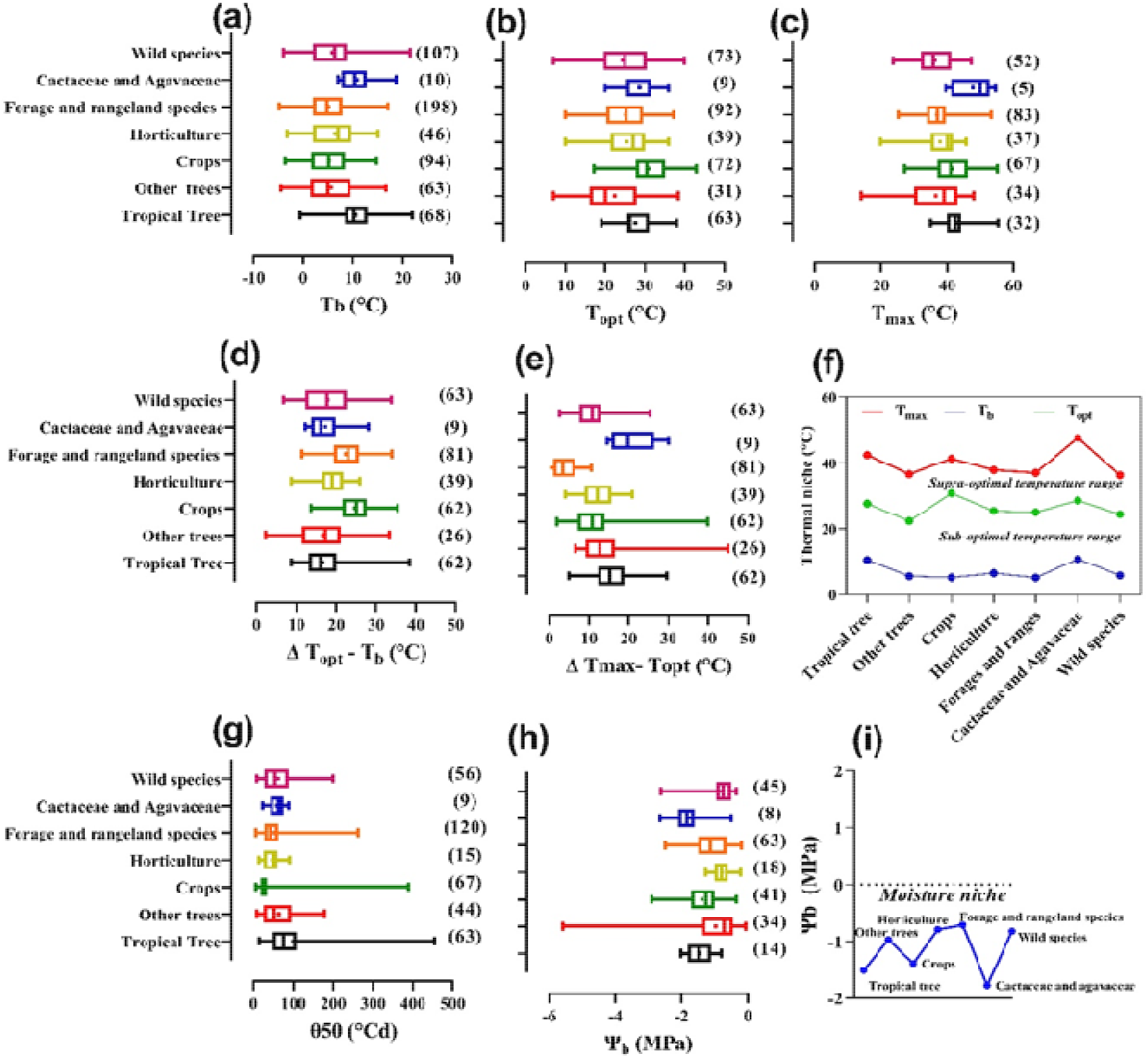
Ranges of base (T_b_; **a**), optimum (T_opt_; **b**), maximum (T_max_; **c**) temperatures, sub-optimal (ΔT_opt_-T_b_; **d**), supra-optimal (ΔT_max_-T_opt_; **e**), thermal niche (upper and lower; **f**), thermal time required to attain 50% germination (_50_; **g**), base water potential (_b_; **h**), and moisture niche (**i**) values of different plant categories considered in this study. The numbers in the parenthesis represent the number of studies found in the literature for each plant group. A frequency distribution was used to show the overview of all traits in some plant categories. In this case, the approach used represents how frequencies are distributed over the observed values. Two-dimensional visualization of the ecological niche of plant species indicating the mean thermal (**f**) and moisture niche (**i**) occupied by each plant category. The breadth of sub-optimal temperature range characterizing the lower thermal niche was estimated by the difference between minimum and optimum temperature while the of supra-optimal temperature range characterizing the upper thermal niche was estimated by the differences between maximum and optimum temperature. Moisture niche was a dimension estimated by taking the mean _b_ values of each plant category from the database.

T_opt_ for germination of crops was limited to a window ranging from 17°C to 44°C, the latter being the highest observed values **(Figure 2b)**. The widest range of T_opt_ for germination was found for wild species, from 7 C to 40 C, followed by the category “other trees” from 6°C to 38°C. In contrast, inter-species variation for T_opt_ for Cactaceae and Agavaceae and tropical trees had the narrowest range, with values of 20 C to 39 C and 20°C to 40°C, respectively.

T_max_ also varied considerably among plant categories. The lowest T_max_ values were observed for non-tropical trees (14C for *Acer saccharum*) and horticultural plants (20°C for *Muscari comosum*). In contrast, the highest T_max_ values were around 55°C, in tropical trees (55.4°C for *Cenostigma microphyllum*), Cactaceae and Agavaceae (54.5°C for *Polaskia chende*), forage and rangeland species (53.3°C for *Urochloa brizantha*) and crops (55°C for *Cicer arietinum L*.) **(Figure 2c)**. Amongst species groupings, T_max_ varied the most for non-tropical trees (14°C for *Acer saccharum* to 48°C for *Anadenanthera colubrine*; **Figure 2c**) and the least for Cactaceae and Agavaceae (from 39.5°C to 54.5°C for *Manfreda brachystachya* and *Polaskia chende*, respectively).

### Thermal niche for germination and _50_

Values of sub- and supra-optimal temperature range varied markedly among plant species (**Figure 2d,e,f**). Forage and rangeland species and crops, showed the widest sub-optimal temperature range, with the extreme values of 20°C and 26°C, respectively. In contrast, non-tropical and tropical trees had the narrowest sub-optimal temperature range, with an average value of 17C for both plant categories. In the supra-optimal temperature range all species had a thermal niche spanning mean values of 12°C to 19°C, e.g. Cactaceae and Agavaceae (19°C), tropical trees (15°C), non-tropical trees (14°C), crops (11°C) and wild species and horticulture species (both 12°C).

_50_ values varied nearly two orders of magnitude among plant groups ranging from 6 °Cd to 500Cd **(Figure 2g)**. Tropical tree species had the highest range of _50_ values, ranging from 15.1°Cd for *Anadenanthera colubrina* to 477°Cd for *Araucaria angustifolia*. In crops, _50_ ranged only about 10-fold, from 6.3°Cd for *Sesamum indicum* L. to 59.4°Cd for *Vicia variabilis*. Wild species, forage and rangeland species, and non-tropical trees showed a similar range in terms of thermal time required for germination, varying from 6°Cd to 263°Cd. Cactaceae and Agavaceae all had relatively short _50_ varying only two-fold from 39.3°Cd to 87.6°Cd.

### Ψ_b_ and moisture niche for germination

Ranges of Ψ_b_ values are presented **Figure 2h**. Compared to the other traits mentioned above, we found fewer data on Ψ_b_ values of tropical trees (14 papers) and horticultural plants (18 papers). In contrast, much more information was available on Ψ_b_ values of forage and rangeland species (63 papers) and wild species (45 papers). Overall, Ψ_b_ values differed considerably among plant species, ranging from a lowest value of -6 MPa for *Atriplex halimus* (a non-tropical tree species) to nearly -0.2 MPa observed for *Stipa grandis* (a forage and rangeland species). Wild species, Cactaceae and Agavaceae, and crops showed similar Ψ_b_ values, with a range varying from -3 MPa to nearly 0 MPa. In contrast, the narrowest range of Ψ_b_ values was for horticultural crops (from -1.27 MPa for *Cucurbita pepo* to -0.21 MPa for *Trachyspermum ammi*) followed by that for tropical trees (from -2.02 MPa for *Apeiba tiborbou* to -0.81 MPa for *Peltophorum dubium*) and wild species (from -2.62 for *Pectocarya heterocarpa* to -0.34 MPa for *Erodium cicutarium*). Forage and rangeland species showed the highest Ψ_b_ values (near to zero as for example -0.21 MPa for *Stipa grandis*).

Values of moisture niche are presented in **Figure 2i**. Similar to the higher thermal niche, tropical trees (14 species and 13 papers), and Cactaceae and Agavaceae (12 species and 8 papers) were revealed to be the most tolerant groups to drought, with a range varying from -2.64 MPa to -0.5 MPa. In contrast, non-tropical trees, and forage and rangeland species showed higher (less negative) Ψ_b_ values, of -0.07 and -0.19 MPa, respectively. Crops had the widest moisture niche, with a range varying from 0 MPa to -1.39 MPa. Forage and rangeland, and horticulture species have the narrowest moisture niche, with the occupancy range varying from 0 to -0.71 MPa and from 0 to -0.79 MPa, respectively.

### Correlations between traits

A positive correlation was found between T_b_ and T_opt_ values for all plant categories (**Figure 3a)**. The steepness of regression slopes was the highest for forage and rangeland species (0.07; P=0.02; **Table 2**) and lowest for Cactaceae and Agavaceae and tropical tree species (0.02 and P=0.00 for both species; **Table 2**).

**Figure 3.**
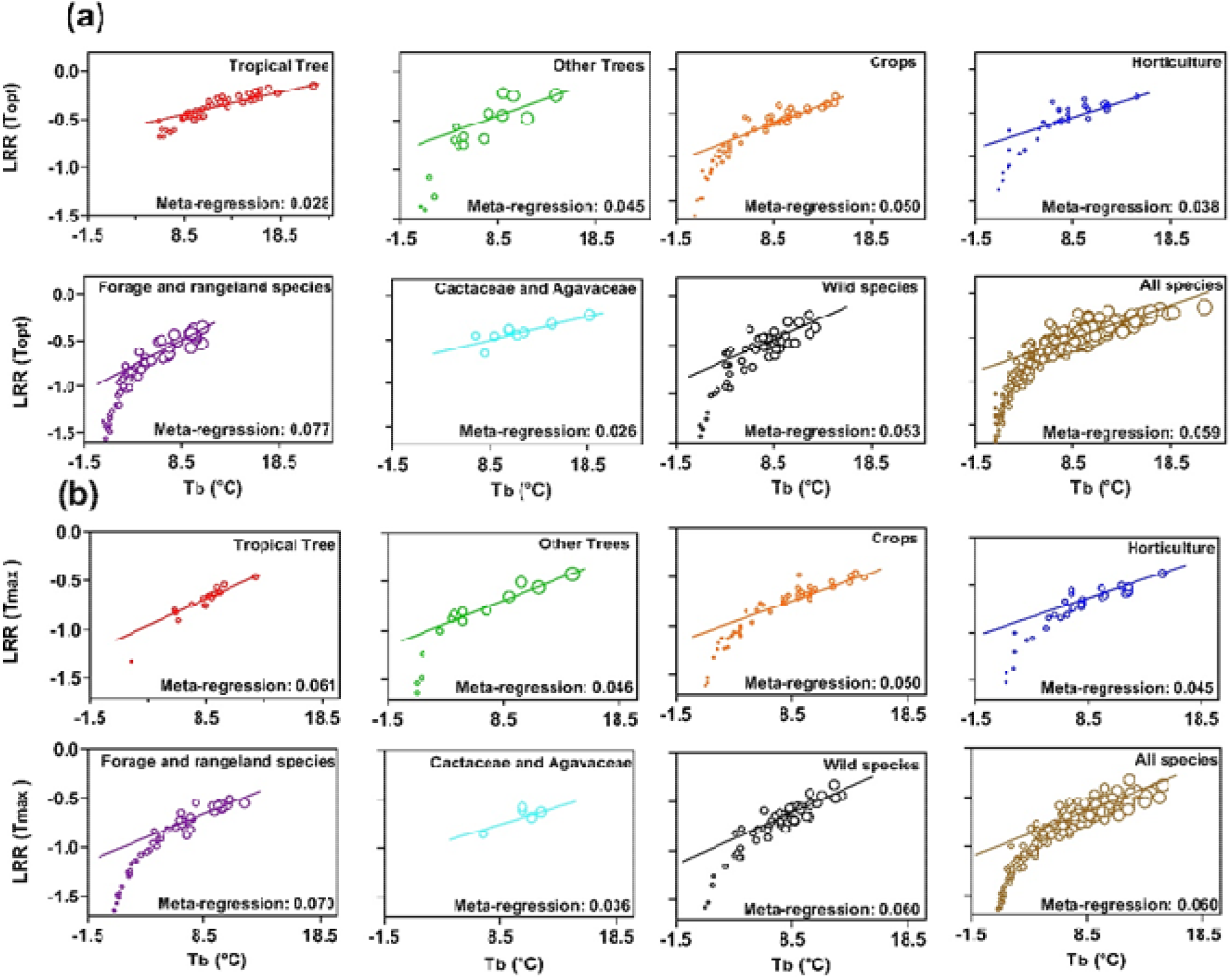
Regression of log risk ratio (LRR) of optimum (a) and T_max_ (b) temperature on base temperature (T_b_). The size of each bubble is inversely correlated with the variance of the log relative risk estimate with larger bubbles showing more inflated variance. LRR represents the probability of changes in the range of T_opt_ and T_max_ as a function of differing T_b_ ranges. Fitted lines were obtained through linear regression approach (*Y=*_*0*_*+X +*). Each plant category is indicated as a separate panel.

**Table 2.** Meta-Regression reporting T_opt_, T_max_, ΔT_opt_ -T_b_, ΔT_max_ –T_opt_, 50 and _b_, values for different categories of plant species as a function of T_b_ for germination. The coefficients were obtained using linear regression. Residual heterogeneity indicates variation around the true regression line. Tau-squared shows the variance of the effect size across studies included in the analysis. T_b_, T_opt_ and T_max_, ΔT_opt_ -T_b_, ΔT_max_–T_opt_ indicate the base, optimum, maximum sub- and supra optimal temperature values, respectively; θ_50_ indicates thermal time required to attain 50% germination while _b_ indicates the base water potential values.

| Plant group | Coefficient |  | SE | P-value | Residual heterogeneity | Tau-squared |
| --- | --- | --- | --- | --- | --- | --- |
|  | Intercept | Slope |  |  |  |  |
| T<sub>opt</sub> |  |  |  |  |  |  |
| Tropical trees | -0.71 | 0.02 | 0.00 | 0.00 | 86.44 | 0.00 |
| Other trees | -0.87 | 0.04 | 0.01 | 0.03 | 22.41 | 0.00 |
| Crops | -1.05 | 0.05 | 0.00 | 0.00 | 89.24 | 0.00 |
| Horticulture | -0.84 | 0.03 | 0.00 | 0.03 | 52.40 | 0.00 |
| Forage and rangeland | -1.17 | 0.07 | 0.00 | 0.02 | 74.14 | 0.00 |
| Cactaceae and Agavaceae | -0.71 | 0.02 | 0.0 | 0.00 | 6.38 | 0.00 |
| Wild species | -0.95 | 0.05 | 0.00 | 0.01 | 10.02 | 0.01 |
| Average | -1.08 | 0.05 | 0.00 | 0.014 | 53.04 | 0.01 |
| T<sub>max</sub> |  |  |  |  |  |  |
| Tropical trees | -1.20 | 0.06 | 0.00 | 0.01 | 14.30 | 0.00 |
| Other trees | -1.07 | 0.04 | 0.00 | 0.01 | 18.68 | 0.00 |
| Crops | -1.16 | 0.05 | 0.00 | 0.00 | 74.76 | 0.00 |
| Horticulture | -1.06 | 0.04 | 0.00 | 0.04 | 45.38 | 0.00 |
| Forage and rangeland | -1.26 | 0.07 | 0.00 | 0.02 | 83.64 | 0.00 |
| Cactaceae and Agavaceae | -1.03 | 0.03 | 0.01 | 0.03 | 4.69 | 0.00 |
| Wild species | -1.15 | 0.06 | 0.00 | 0.02 | 53.17 | 0.00 |
| Average | -1.23 | 0.06 | 0.00 | 0.00 | 55.31 | 0.01 |
| ΔT<sub>opt</sub>-T<sub>b</sub> |  |  |  |  |  |  |
| Tropical trees | -0.84 | 0.06 | 0.00 | 0.04 | 51.12 | 0.00 |
| Other trees | -0.69 | 0.04 | 0.00 | 0.00 | 17.58 | 0.00 |
| Crops | -1.02 | 0.06 | 0.00 | 0.03 | 12.46 | 0.01 |
| Horticulture | -0.87 | 0.05 | 0.00 | 0.01 | 52.93 | 0.00 |
| Forage and rangeland | -1.19 | 0.09 | 0.00 | 0.00 | 62.39 | 0.00 |
| Cactaceae and Agavaceae | -0.74 | 0.04 | 0.00 | 0.02 | 8.11 | 0.00 |
| Wild species | -0.97 | 0.07 | 0.00 | 0.02 | 82.97 | 0.01 |
| Average | -1.01 | 0.07 | 0.00 | 0.00 | 63.75 | 0.01 |
| $\Delta T_{\max} - T_{\text{opt}}$ | | | | | | |
| Tropical trees | -0.938 | 0.07 | 0.00 | 0.04 | 31.16 | 0.00 |
| Other trees | -1.15 | 0.09 | 0.00 | 0.00 | 27.32 | 0.01 |
| Crops | -0.79 | 0.06 | 0.00 | 0.03 | 17.46 | 0.01 |
| Horticulture | -0.71 | 0.06 | 0.00 | 0.01 | 22.53 | 0.00 |
| Forage and rangeland | -0.91 | 0.07 | 0.00 | 0.00 | 12.25 | 0.00 |
| Cactaceae and Agavaceae | -1.03 | 0.07 | 0.00 | 0.02 | 18.56 | 0.01 |
| Wild species | -0.79 | 0.08 | 0.00 | 0.02 | 62.67 | 0.01 |
| Average | -0.65 | 0.05 | 0.00 | 0.00 | 52.25 | 0.00 |
| 50 |  |  |  |  |  |  |
| Tropical trees | 1.25 | -0.03 | 0.01 | 0.02 | 60.32 | 0.06 |
| Other trees | 1.02 | -0.03 | 0.01 | 0.00 | 63.74 | 0.09 |
| Crops | 0.52 | -0.04 | 0.03 | 0.01 | 30.18 | 1.18 |
| Horticulture | 1.61 | -0.11 | 0.02 | 0.03 | 11.79 | 0.07 |
| Forage and rangeland | 0.82 | -0.02 | 0.05 | 0.02 | 19.38 | 0.00 |
| Cactaceae and Agavaceae | 0.93 | -0.02 | 0.02 | 0.00 | 8.48 | 0.00 |
| Wild species | 1.40 | -0.07 | 0.00 | 0.00 | 63.02 | 0.01 |
| Average | 0.90 | -0.02 | 0.00 | 0.01 | 21.50 | 0.12 |
| b |  |  |  |  |  |  |
| Tropical trees | 0.08 | 0.40 | 0.41 | 0.03 | 0.33 | 0.00 |
| Other trees | -1.38 | -0.44 | 0.22 | 0.02 | 8.67 | 0.00 |
| Crops | -1.30 | -0.60 | 0.13 | 0.01 | 18.20 | 0.08 |
| Horticulture | -1.12 | -0.38 | 0.37 | 0.01 | 19.61 | 0.01 |
| Forage and rangeland | -0.50 | -0.09 | 0.04 | 0.01 | 55.63 | 0.03 |
| Cactaceae and Agavaceae | -1.70 | -0.54 | 0.30 | 0.02 | 1.60 | 0.00 |
| Wild species | -1.27 | -0.73 | 0.14 | 0.01 | 40.08 | 0.01 |
| Average | -0.003 | -0.26 | 0.05 | 0.00 | 30.29 | 0.05 |

Values of T_b_ positively correlated with T_max_, with a high proportion of variance corresponding to high T_b_ values (**Figure 3b)**. The steepness of the regression slopes differed markedly among plant categories, from 0.07 for Forage and Rangeland species (P=0.02 and 0.01, respectively; **Table 2**), then wild species and tropical trees, (0.06 for both, P=0.02 and 0.00, respectively; **Table 2**). In contrast, a weaker correlation was observed between T_b_ and T_max_ in Cactaceae and Agavaceae, and horticulture species as shown by the regression slope values of 0.03 and 0.04, respectively (P=0.03 and 0.04, respectively; **Table 2**).

Different data inputs and variance are shown in **Figure 3** by varying size of bubbles such that inflated bubbles refer to higher variance. Despite differing levels of data coverage that may influence the slopes of meta-regression and subsequently the correlations obtained, the general trend indicates that there is a positive correlation between T_b_ and T_max_ (the slope of 0.060; **Table 2; Fig 3b**). Tropical trees, and forage and rangeland species showed steeper slopes than the general trend while all other plant categories were characterized by values lower than the general trend (**Table 2; Fig 3b**). Overall, variance inflated with increasing T_b_, indicating that theres is considerable difference between species groups and data shortage **(Figure 3)**. This trend may be reflected in other parameters as well.

Correlation results between T_b_ and thermal niche are presented in **Figure 4**. We found a positive correlation between T_b_ and ΔT_opt_ -T_b_ values for all plant categories **(Fig 4a)**. The steepness of regression slopes was the highest for forage and rangeland species (0.095; P=0.00; **Table 2**), followed by that of wild (0.079; P=0.02; **Table 2**), crops (0.063; P=0.03; **Table 2**), tropical tree (0.061; P=0.04 Table 1), horticulture (0.0596; P=0.01 Table 1), Cactaceae and Agavaceae (0.048; P=0.02), and non-tropical tree (0.04; P=0.00; **Table 2**).

**Figure 4.**
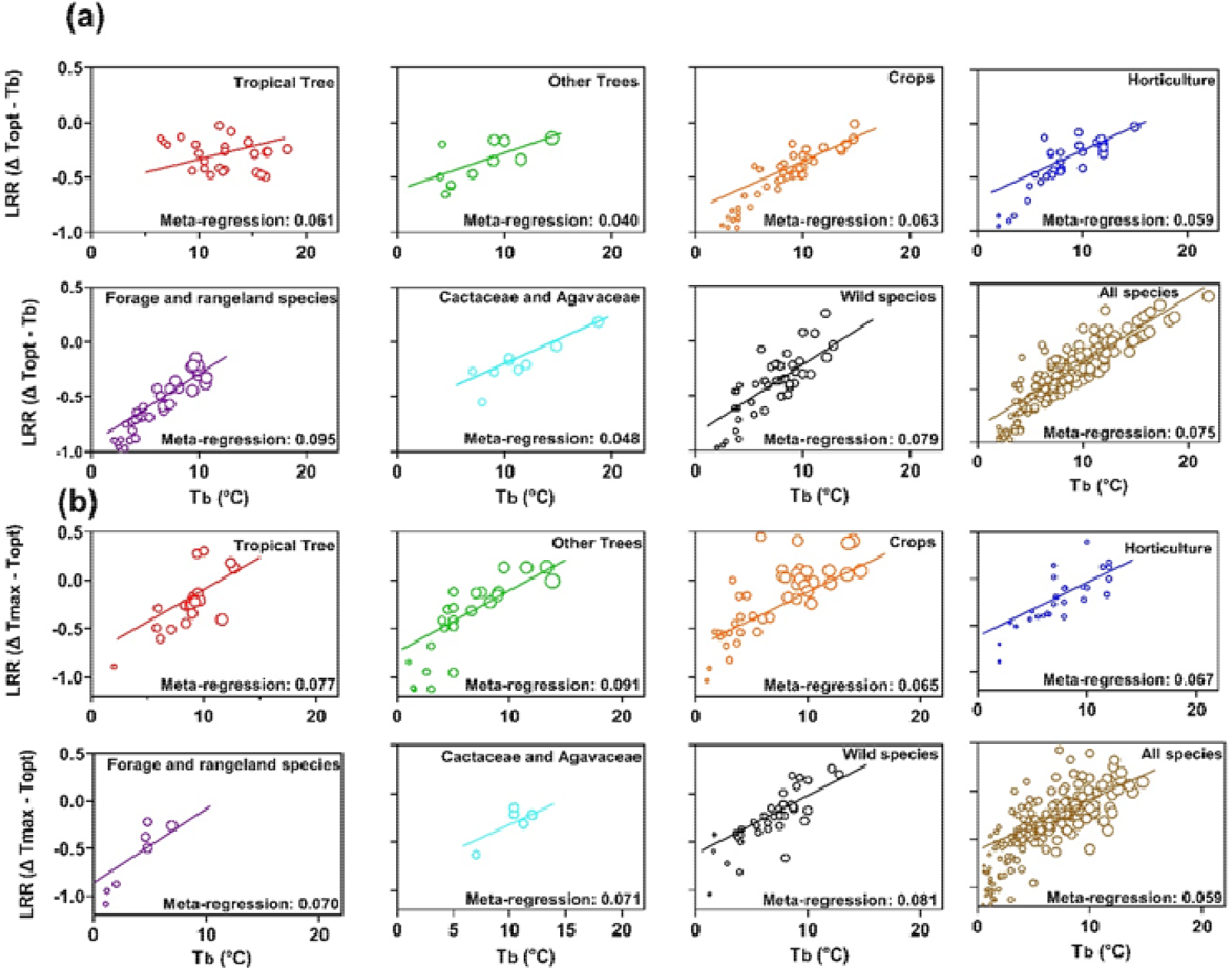
Regression of log risk ratio (LRR) of sub-optimal (ΔT_opt_-T_b_; **a**), supra-optimal (ΔT_max_-T_opt_; **b**), on base temperature (T_b_). The size of each bubble is inversely correlated with the variance of the log relative risk estimate with larger bubbles showing more inflated variance. LRR represents the probability of changes in the range of Δ values of sub- and supra-optimal temperature as a function of changes in T_b_. Fitted lines were obtained through linear regression approach (*Y=*_*0*_*+X +*). Each plant category is indicated as a separate panel.

There was also a positive correlation between T_b_ and ΔT_max_ –T_opt_ values for all plant categories (**Fig 4b**). Other trees and wild species showed the highest steepness of regression slopes, with the slope of 0.091 and 0.081(P=0.00; **Table 2**). Followed by other trees (0.091) and wild species (0.081), the highest slope was for tropical trees (0.077) and forage and rangeland species (0.070). In contrast, the steepness of regression slopes was the lowest for crops (0.065).

Correlation results between T_b_ and _50_ are presented in **Figure 5a**. We observed a negative relationship between T_b_ and _50_ with different meta-regression values among plant categories. The steepness of slope was the highest in horticultural species (i.e. -0.11; **Table 2**) followed by wild species (- 0.07) while the lowest slope was observed for crops (- 0.04). Horticultural species were followed by Cactaceae and Agavaceae, and forage and rangeland species showed a slight increase in _50_ as a function of T_b_, with the slope steepness of -0.020 and - 0.022, respectively **(Table 2)**.

**Figure 5.**
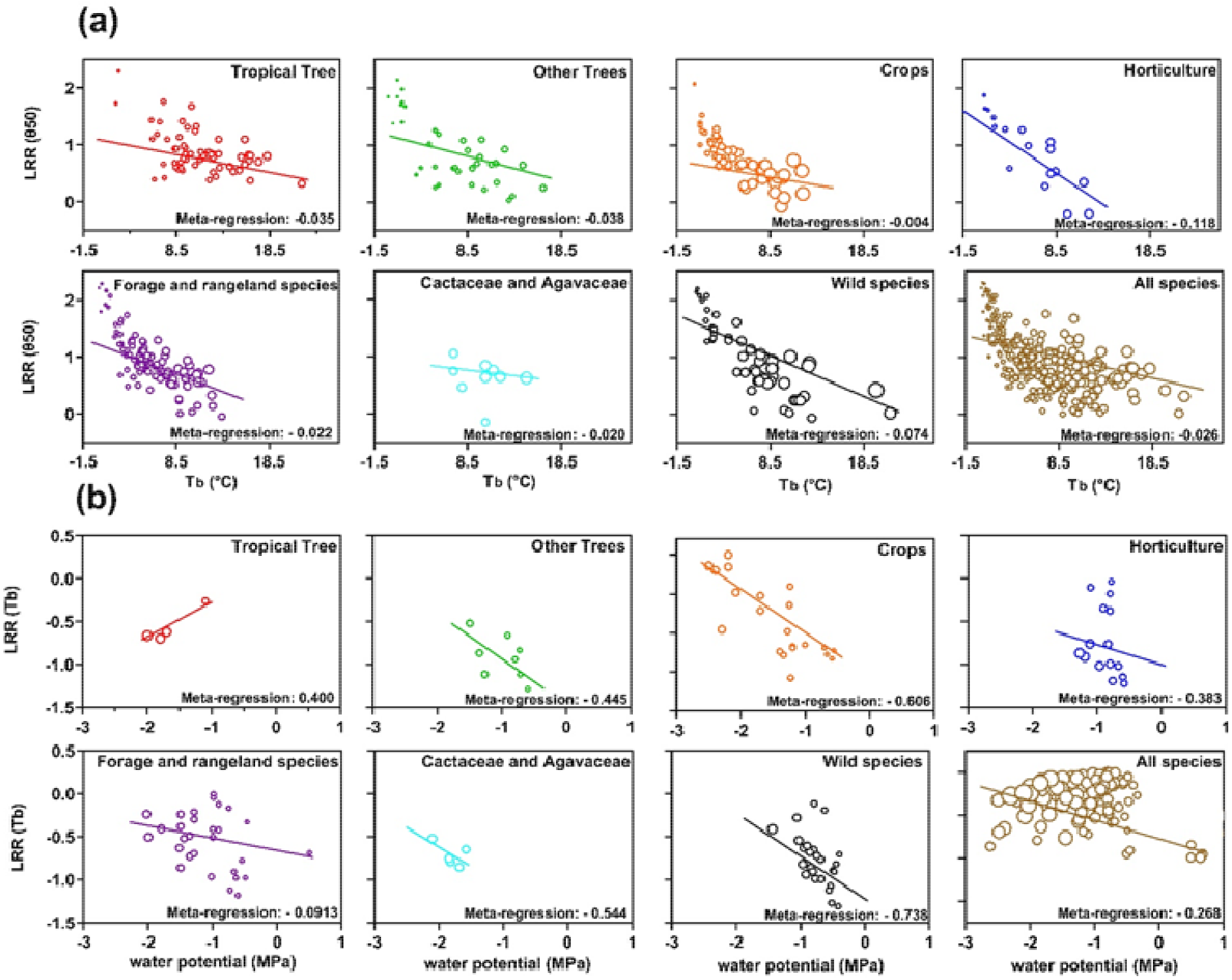
Regression of log risk ratio (LRR) of thermal time required to attain 50% germination (θ50%) on base temperature (T_b_; **a**) and LRR of base temperature (T_b_) on base water potential (_b_) values (**b**). The size of each bubble is inversely correlated with the variance of the log relative risk estimate with larger bubbles showing more inflated variance. LRR represents the probability of changes in θ50% as a function of changes in T_b_ **(a)** values and in the range of T_b_ as a function of differing _b_ values **(b)**. Fitted lines were obtained through linear regression approach (*Y=*_*0*_*+X +*). Each plant category is indicated as a separate panel.

Overall, a negative relationship was found between T_b_ and Ψ_b_ values in all species included in this study, with the regression slope of -0.26 (−0.26 P=0.00; **Table 2**). A strong negative correlation between T_b_ and _b_ values was observed for all but tropical tree species (**Figure 5b)**, with considerable difference in the steepness of regression slopes among plant categories (P=0.03; **Table 2**). Wild species had the lowest values of the regression slopes (−0.73; P=0.01; Table 1), followed by those of crops (−0.60; P=0.01), Cactaceae and Agavaceae (−0.54; P=0.02; Table 1), non-tropical tress (−0.44; P=0.02; **Table 2**), and horticulture species (−0.38; P=0.01; **Table 2**). In contrast, a positive correlation between the T_b_ and _b_ values was observed for tropical tree species with a slope of 0.40, although this plant category had few data (only 4 species).

## 4. Discussion

Sensitivity to environmental factors is a critical ecological process for plants occurring in unpredictable habitats where successful germination is highly dependent on the spatio-temporal variability of temperature and rainfall (Batlla & Benech-Arnold, 2015; Soltani *et al*., 2017a,b; Maleki *et al*., 2021). For 569 species of wild and cultivated origin, we analysed the thermal and moisture germination niches through temperature and water potential thresholds and thermal timing, and considered how these varied among different species and plant categories.

Thermal germination niche was classified into two distinct ranges viz. sub- and supra-optimal ranges. Within sub-optimal range, crops showed the widest thermal germination niche that could be a result of the selection and domestication process. A wide sub-optimal temperature range increases the possibility of crop adaptations across contrasted thermal environments, including fluctuating temperatures, which may allow the relative synchronization of germination under known environmental and management conditions. The availability of fit-for-purpose seed lots greatly reduces risks of germination failure, particularly at sub-optimal temperatures. Predictable performance at low temperature means that spring crops can be sown early to escape late season summer drought and winter crops sown to avoid early/late season frosts. In contrast to crops with the widest thermal germination niche, species originating from warmer environments, such as tropical trees, had the narrowest thermal niche within the sub-optimal temperature range. This could be due to a strong natural selection and environmental drivers such that thermal niche is narrow when environmental filters are stronger (Fernández-Pascual *et al*., 2017). Consequently, when considering germination niche, habitat-specific strategies should be taken into account as habitat-generalist and habitat-specialist plants can have broader and narrower germination niches, respectively (Marques *et al*., 2014). Although we did not focus on habitabt-specific strategies, our results highlight the need for future research on the environmental and abiotic conditions (i.e., environmental and physico-chemical) which may limit the emergence of plant species (Tudela-Isanta *et al*., 2018). We also show that crops relative to tropical trees have the steepest regression slope when considering the relationship between T_b_ and T_opt_ and delta values, indicating that crops have higher sensitivity to sub-optimal temperatures compared to tropical trees (**Figure 4a; 5a**). Such enhanced sensitivity effectively means that crop seed germination is a more efficient over a range of conditions.

In contrast to sub-optimal temperature, the sensitivity pattern of supra-optimal temperature range was considerably different, with Cactaceae and Agavaceae having the widest and crops showing the narrowest supra-optimal temperature range (**Figure 2f**). This suggests that exposure of Cactaceae and Agavaceae to extreme tropical dryland conditions may have facilitated population divergence through an adjustment in germination behaviour as a consequence of parental environmental (Lampei *et al*., 2017). Based on the concept of parental environmental effects, offspring can take advantage of information about parental environment to drive specific phenotypes for optimizing the match with the offspring environment (Lacey, 1988). This ecological process requires a correlation between parental and offspring environments (Ezard *et al*., 2014; Burgess & Marshall, 2014; Leimar & McNamara, 2015). In contrast to Cactaceae and Agavaceae, crops, which are often grown under optimal management conditions, have a lower capacity to tolerate warmer conditions, and thus, could be subjected to higher mortality risks under climate change. An example is the reduced capabity of sunflower (*Helianthus annuu*s) compared to its wild relatives to convert to normal seedlings at high temperatures and low water potentials (Castillo-Lorenzo *et al*., 2019a,b) Our meta-regression results highlight a sensitivity pattern whereby, within both sub- and supra-optimal temperature ranges, species (or gropus) with lower T_b_ might be the most sensitive to climate change because of their smaller delta values (i.e., T_o_-T_b_; T_max_ – T_o_). This means that increases in global average temperature could impact such species germination performance as the supra-optimal temperature range would be relatively easier to reach. Most likely, and if genotypic limits are not exceeded, natural selection for and evolution towards higher T_b_ and wider delta values could enable continuing emergence. Such adaptation to a potentially new (but still narrower) germination niche (Fernández-Pascual *et al*. (2017) might be more likely for annual species than pererennials, such as trees, Cactaceae and Agavaceae.

The trend of supra-optimal temperature range also reflects the moisture niche dependency, with Cactaceae and Agavaceae occupying broader moisture niche. Again, this may suggest that divergence in parental environmental effects could have driven Cactaceae and Agavaceae toward wider moisture niche to be able to capitalize on short periods of water availability under unpredictable conditions (Baskin & Baskin, 2014; Lampei *et al*., 2017) and the rapid initiation of the germination process as soil moisture increases. Overall, we show that species with narrower moisture niche have a stronger relationship between T_b_ and Ψ_b_ explaining that species with wider supra-optimal temperature range may have broader moisture niche, which results from a correlation between parental and offspring environments (Ezard *et al*., 2014; Burgess & Marshall, 2014; Leimar & McNamara, 2015).

The sensitivity pattern reflected in sub- and supra-optima temperature range is also evidence in the Ψ_b_ values. Species with higher T_b_ are potentially more tolerant to drought as shown by lower Ψ_b_ values. For instance, tropical trees show a positive correlation between T_b_ and Ψ_b_ that is probably due to a high frequency of rainfall in tropical areas. However, the ongoing variability in rainfall frequency and intensity due to climate change may affect tropical trees through the germination phase of the life cycle (Pritchard *et al*., 2022).

### Variability in T_b_ values among plant categories

Cactaceae and Agavaceae are characterized by the lowest variabilities in terms of T_b_ extreme values, suggesting their sensitivity to lower temperature range. A negative correlation was observed between T_b_ and _50_ values. Annual plant species of tropical origin (e.g. cotton and mungbean), have the highest T_b_ values, while tree species from cooler regions (e.g. such as oak, betula and ash) show the lowest T_b_ values. Moreover, some crops (e.g. winter pea) are projected to be able to germinate at sub-zero temperatures (T_b_ = -1.10 °C), which could be due to selection through breeding (Stupnikova *et al*., 2006) Overall, we found positive correlations between T_b_ and other traits related to cardinal temperatures suggesting that plant species alter T_b_ values in harmony with other thermal traits as a highly efficient adaptation strategy to coping with harsh conditions.

Based on the results of this study, T_b_ for some plant categories are below zero except for Cactaceae and Agavaceae and tropical trees. Germination at subzero temperatures is likely because proteins and sugars in plant cell that protect cell environment from damages caused by ice formation. A recent study (Jaganathan *et al*., 2020) proposed three main mechanisms for sustaining survival at subzero temperatures, including the existence of water impermeable seed coats, the super-cooling of seed tissue and freezing tolerance triggered by extracellular-freezing. Species included in our study might take advantage of these mechanisms to ensure their survival at subzero temperatures.

### Amplitude of T_opt_ and T_max_ values among plant categories

The range of T_opt_ was narrow compared to T_max_ range. The range of T_max_ ranged from 8C in forages and rangeland species to 44C in crops, suggesting that crops seem to be a category that has higher T_opt_ (from 17.2C for *Onobrychis subnitens* to 43C for *Sorghum bicolor*) due to domestication and selection events targeting rapid germination across fluctuating environmental conditions.

Crops and horticultural species are likely to have higher T_opt_ and T_max_ values compared with other categories. This could be driven by their selection aimed at adapting to wider environmental conditions, including hotter regions of the world. For instance, horticultural species are high-value crops that are most often grown under irrigated conditions that make it possible to grow them even across extremely hot conditions. Crops with tropical origin, such as cotton and mungbean, have the highest T_opt_ and T_max_ values. For many species, the range of T_max_ was 40-45 C, and it seems that seeds of these species are not able to progress towards completion of germination above T_max_. However, some crops are able to germinate at temperatures> 45°C (e.g. *Phaseolus vugaris* L., *Glycine max* L., *Cicer arietinum* L. and *Sesamum indicum* L.). These values suggest that there may be an upper temperature limit (T_max_) to the germination process around 50-55°C. This limit could be imposed by the onset of enzyme denaturation and activities at molecular level leading to cell death and failure of cell growth. For example, lentil and pea seed amylase required for starch degradation in the early stages of germination becomes inactive within minutes of exposure to 70°C and 80 °C, respectively (Tárrago & Nicolás, 1976; Adegbanke *et al*., 2021).

This suggests that in addition to adaptation to environmental conditions within genetic limits, there might be cell functional and signaling constraints that control the biological range within which seeds are able to germinate. In contrast, the threshold-type response of wild species (other than Cactaceae and Agavaceae) to T_max_ was weaker than that of crops and was restricted to 20-45°C. For instance, non-tropical trees such as sugar maple (*Acer saccharum* M.) have the lowest T_max_ values (i.e. 14°C). Such a low T_max_ value may be confounded by the presence of dormancy in this species. Species with such non-deep physiological dormancy have limited cardinal temperatures at maturity, e.g. *Aesculus hippocastanum* (Pritchard et al., 1999), and seeds with this ‘conditional dormancy’ progressively gain the ability to germination over a broader temperature range (Baskin & Baskin, 2014; Maleki *et al*., 2021; Soltani *et al*., 2022). As for T_opt_ values, they increased as a function of T_b_, as shown by the regression slope, which may suggest coevolution of threshold-type responses to temperature that need to be explored by future research.

### Range of _50_ values among plant categories

Values of _50_ provide a context for adaptive strategies optimizing the efficiency of germination in relation to temperature and impacting timing (Donohue *et al*., 2010; Maleki *et al*., 2021). Higher thermal requirement is mainly linked to dormancy status (Baskin & Baskin, 2014; Maleki *et al*., 2021), although dormancy-breaking treatment may be applied to overcome seed dormancy. Tropical trees, non-tropical trees and wild species are shown to have longer _50_ (i.e. larger thermal inputs for the same germination proportion). This could be a valuable adaptation to cope with changing environments, such that germination is spread and the likelihood of all seeds in a cohort dying due to unfavorable conditions is reduced. In contrast, crops showed the smallest _50_ values, suggesting domestication has resulted in selection for fast germination and high vigour. This is also true for horticultural species, particularly ornamental or medicinal species, that have been selected for a specific use (provisioning, nutrients) and cropping cycle. A similar narrow range of short thermal requirements (i. e. lowest _50_ values) for germination found in horticultural species and crops is evident in Cactaceae and Agavaceae. Rather than domestication and selection being important, for the latter groupings an adaptive germination strategy in response to harsh environments is likely. In this way, Cactaceae and Agavaceae can take advantage of temporary suitable conditions and rapidly complete this critical step (germination) in their life cycle. Such considerations support the hypothesis of a bet-hedging strategy for germination that depends on specific environmental conditions (Gremer *et al*., 2016).

### Extent of _b_ values among plant categories

Unlike for traits related to T_b, 50_ and cardinal temperatures, we found less literature on Ψ_b_ with only for 226 species compared to T_b_ and T_opt_ reported for 461 and 334 species, respectively. Nonetheless, studies published in the last decade have focused more on seed Ψ_b_ compared with those published before 2011 (database of Dürr *et al*., 2015). Undoubtedly, with concerns growing about the wider impacts of climate change on drought, a wider dataset on species’ germination responses to water potential would be valuable.

Overall, the lowest values of Ψ_b_ were found for wild woody species, in particular -5.58 MPa for *Atriplex halimus*. This shrub species is adapted to extreme environments, being highly drought resistant and tolerant of saline conditions, and highlights one of the most important aspects of our new data: to identify model species at the limits of environmental adaptability for further study.

Although we found 65 and 45 studies reporting Ψ_b_ values of forage and rangeland species, and wild species, respectively, the dearth of information on tropical trees and horticultural plants (14 and papers, respectively) calls for further research as germination response to water availability is one of the most important environmental driver for adaptation of plant species.

### Contrasted germination niches among plant categories

We found that sensitivity to environmental factors is a critical ecological process for plants occurring in unpredictable habitats where successful germination is highly dependent on the spatio-temporal variability of temperature. Therefore, germination niche, as a fundamental ecological indicator, should be included more in future risk assessments to species survival. Environmental factors may determine germination niche through regulating both seed dormancy and germination behaviour (Batlla & Benech-Arnold, 2015; Soltani *et al*., 2017a,b; Maleki *et al*., 2021). We found distinct thermal and moisture germination niche, which is defined as limits characterizing species distribution, for various groups of species. Moreover, differential thermal times for germination describes changes in the sensitivity of germination responses to randomly changing environments, leading to various strategies adopted by plants in term of niche construction and regulation of germination process. Maternal thermal environments may play a regulatory role in niche construction. *Cactaceae* and *Agavaceae*, for example, have the widest Supra-optimal temperature range, suggesting that species experiencing higher temperatures during their lifetime, or more precisely species having experience of higher temperatures, should have wider thermal niche, which is consistent with the ecological concept of germination niche, in which species construct their niche in response to environmental conditions they have experienced (Sultan, 2015; Fernández-Pascual *et al*., 2019). Although *Cactaceae* and *Agavaceae* have wider upper thermal niche, a recent study showed that these plants are sensitive to increasing global temperatures as the thermal environments in which they inhabit are close to their upper thermal limit (Sentinella *et al*., 2020). Similarly, species facing dry environments could be generally more tolerant to drought condition by having wider moisture niche as seen in *Cactaceae* and *Agavaceae*. However, in some cases, species from wetter environmental conditions may be better tolerant to drought as observed in *Brassica* sp. (Castillo-Lorenzo *et al*., 2019a). Interestingly, plant groups manipulated to adapt to changing environmental conditions, such as crops and horticultural species, are better suited to lower temperatures by showing wider lower thermal niche. However, these plants are highly sensitive to higher temperatures as shown by a narrow upper thermal niche, suggesting that genetic manipulation and agronomic practices applied to these plant species (e.g. irrigation) may have led to these changes.

## 5. Conclusion

In this study, we extracted data from 377 studies conducted worldwide on 569 plant species including trees, grasses, crops and wild species to determine ranges of threshold-type responses to temperature and water potential, as two main environmental filters, to determine relationship among these traits. Despite all the complexities in collecting and interpreting data, our results show a strong relationship among ecologically meaningful traits quantifying ranges of threshold-type responses to temperature and water potential and the results of our meta-regression provide insight into the adaptation of various categories of species distributed worldwide to the environments in which they inhabit, factors constraining adaptation and the role of domestication under current and future climate. We found that, under environmental disturbances, distinct germination traits expressed among and within species can allow ecosystems to persist since such adaptive traits are likely to determine threshold-type values for life-history transitions, particularly germination as an important determinant of plant recruitment. Because climate change is expected to become increasingly severe in the future, any changes in threshold-type responses to temperature and water potential may considerably influence plant function and its performance. A better understanding of functional traits and precise field-based validation underlying adaptation would be useful in providing detailed information on plant regeneration.

## Author contributions

JRL conceived the original idea. ES collected and extracted the data from the literature. KM assembled and managed the database, analyzed the data and drafted the manuscript. JRL and ES supervised the work. KM, JRL, ES, CES and HWP commented on the data analyses and revised the manuscript. All authors approved the final version.

## Acknowledgments

The authors thank Carolyne Dürr for her earlier feedback on this work. JRL received financial support from the AgroEcoSystem division of INRAE.

## Data availability

The dataset has been deposited to a data repository of INRAE (Data identification number: XP3XHW_2022). A version of record of the repository can be found at https://doi.org/10.15454/XP3XHW.

